# Unravelling the mystery of endemic versus translocated populations of the endangered Australian lungfish (*Neoceratodus forsteri*)

**DOI:** 10.1101/2023.03.24.533984

**Authors:** Roberto Biello, Silvia Ghirotto, Daniel J. Schmidt, Silvia Fuselli, David T. Roberts, Tom Espinoza, Jane M. Hughes, Giorgio Bertorelle

## Abstract

The Australian lungfish is a primitive and endangered representative of the subclass Dipnoi. The distribution of this species is limited to south-east Queensland, with some populations considered endemic and others possibly descending from translocations in the late nineteenth century shortly after European discovery. Attempts to resolve the historical distribution for this species have met with conflicting results based on descriptive genetic studies. Understanding if all populations are endemic or some are the result of, or influenced by, translocation events, has implications for conservation management. In this work, we analysed the genetic variation at three types of markers (mtDNA genomes, 11 STRs, and 5,196 nuclear SNPs) using the Approximate Bayesian Computation algorithm to compare several demographic models. We postulated different contributions of Mary River and Burnett River gene pools into the Brisbane River and North Pine River populations, related to documented translocation events. We ran the analysis for each marker separately, and we also estimated the posterior probabilities of the models combining the markers. Nuclear SNPs have the highest power to correctly identify the true model amongst the simulated datasets (where the model was known), but different markers typically provided similar answers. The most supported demographic model able to explain the real dataset implies that an endemic gene pool is present in the Brisbane and North Pine Rivers where past translocations are documented. These results will inform ongoing conservation efforts for this endangered and iconic species and warrant careful consideration of future genetic management of the Australian lungfish populations.

## INTRODUCTION

Approximately 120 years ago, a retired bank manager, Daniel O’Connor, addressed the members of the Royal Society of Queensland in Australia: “I have the pleasure to inform you that the work of procuring *Ceratodus* and transferring live specimens to new habitats which you entrusted to me is finished” (O’Connor, 1897). The *“Ceratodus”* genus referred to by O’Connor was the Australian lungfish now known as *Neoceratodus fosteri.* It had only just been discovered in the Burnett River in 1870 and was described a few years later by the Brisbane Museum director as “one of the most famous things in Australia’’ (Longman, 1925). At this point in time, Australian lungfish had not been recorded or documented in any rivers south of its established range in the Burnett and Mary River catchments. The translocations were thus commissioned in response to a perceived risk the species could face extinction on account of its restricted distribution, and the rarity of juveniles observed in the wild.

The Australian lungfish is still present in the Burnett and Mary catchments and in most of those rivers that received documented translocations, although only abundant in a few (Kind, 2011). The contributions of O’Connor in preventing the perceived extinction of the last of at least 11 lungfish species distributed in the Cretaceous all over Australia (Kemp, 1997) remains unclear, with ongoing conjecture as to the endemic distribution of the species prior to translocations (Kemp & Huynen, 2014). Nevertheless, this IUCN endangered species is currently threatened by habitat modifications mainly due to water resource development and land use that reduce the quality and availability of important riverine habitats used for reproduction (Kemp, 2018; Brooks et al., 2019). Conservation efforts and water resource management must find a balance to sustain this threatened species.

The impact of O’Connor’s translocations requires exploration to better prioritize future conservation efforts. Could these early conservation efforts have resulted in backup populations outside of its endemic distribution, or have these efforts created a new genetically mixed population containing a diverse genetic lineage of endemic and introduced gene pools. Lungfish are not only unique amongst Australian fauna, but also a crucial biological and evolutionary model for studying and understanding the evolution of tetrapods and adaptations to land (e.g., Woltering et al., 2020; Meyer et al., 2021). As such, understanding and preventing extinction is a great ethical obligation to the world (Antonelli & Perrigo, 2018).

The main question explored by this research, is whether the translocations that occurred in 1896 were the inception of the present-day lungfish populations in the Brisbane and North Pine Rivers, or they already existed in those rivers and were genetically influenced by those translocations (see **Figure 1**). Fisheries management agencies in Australia consider the Brisbane River and North Pine River populations as introduced (Bishop et al., 2018). The historical translocations described 109 adults being sourced from the Mary River (see **Figure 1**). Of these only five and eight individuals were introduced into the Brisbane River and the North Pine River basins, respectively (O’Connor, 1897). All other survivors were released into other rivers across the South East Queensland region and have largely been unsuccessful in establishing viable populations. Anecdotal evidence suggests translocations may have also occurred later, possibly with individuals sourced from the Burnett River, as mentioned in newspapers from the time. Furthermore, recent information suggests that additional illegal translocations of lungfish may still be occurring (Kind, 2011).

**Figure 1.**
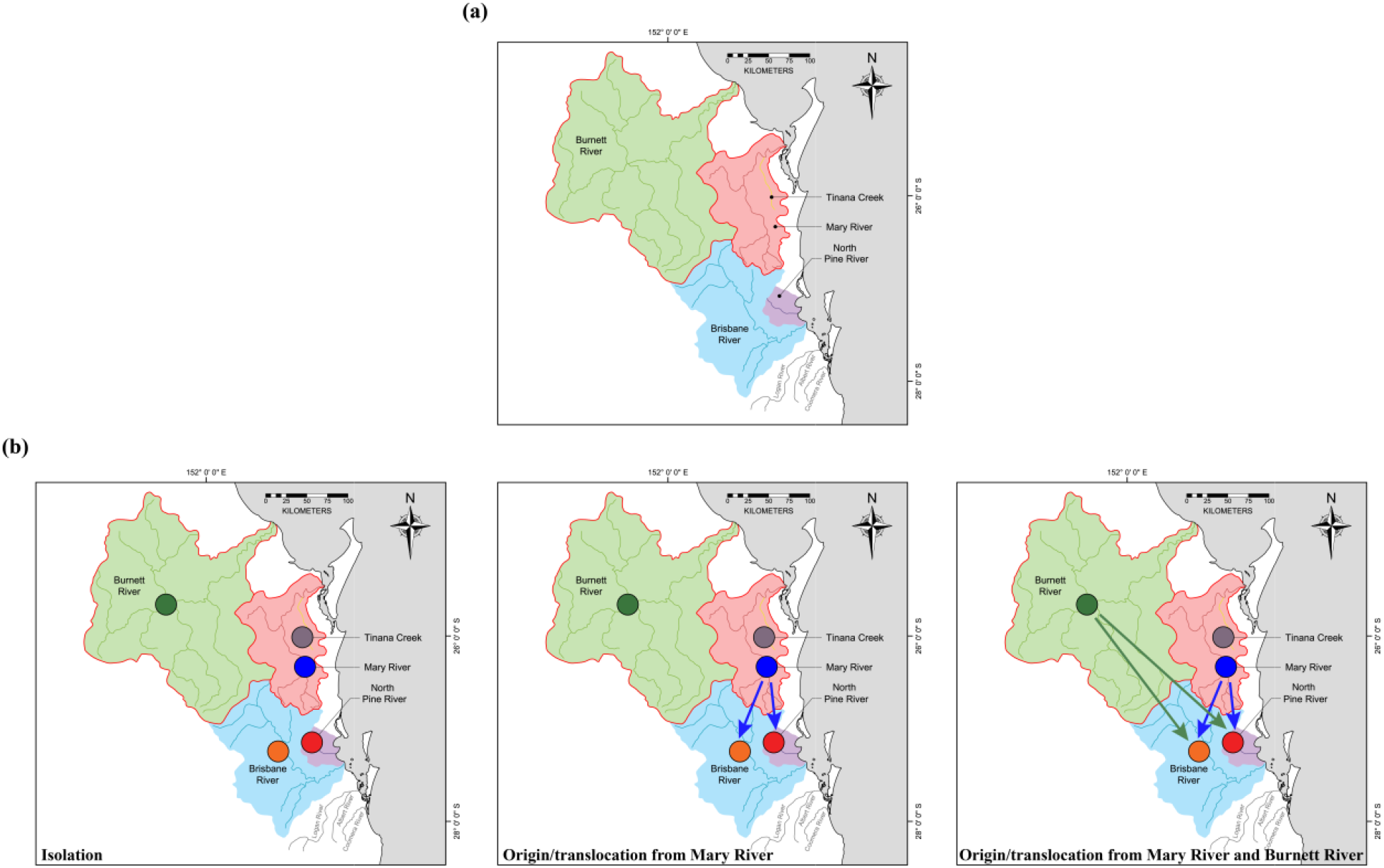
a) Catchment areas of the individuals included in this study. b) Main demographic models tested with ABC-rf; left panel: populations are evolving in isolation since divergence, corresponding to models 1a to 7a in **Figure S1**; central panel: Brisbane and North Pine River populations descend completely (Models 1b to 7b in **Figure S1**) or partially (Models 1d to 7d in **Figure S1**) from individuals translocated from Mary River; right panel: Brisbane River and North Pine River populations descend completely (Models 1c to 7c in **Figure S1**) or partially (Models 1e to 7e in **Figure S1**) from individual translocated from Mary River and Burnett River.

The intention of the original translocation event was to establish new sustainable populations in suitable rivers where the lungfish was not formerly found. Despite many early explorers, naturalists and ichthyologists searching for *Ceratodus* and other unique river fauna within Queensland, there are no historical records describing the occurrence of lungfish in either the Brisbane or North Pine Rivers before the translocations. Diaries of explorers from the 1820’s, who extensively explored the Brisbane and surrounding rivers, sourcing fish for consumption and trading fish with the indigenous, made no mention of a fish unlike those more readily recognisable species (Steele, 1972). Of course, absence of evidence is not evidence of absence, especially for a species with a cryptic habit, a restricted geographical range and not previously described by science. The hypothesis that the current Brisbane River and North Pine River populations descended entirely from the O’Connor translocations has been questioned before (Kemp, 1986). Previous conclusions were often drawn in the absence of biological traits about the species, such as fecundity, longevity, survival rates and key drivers for recruitment, all essential to make informed conclusions about potential population dynamics. The presence of lungfish inhabiting the Brisbane River prior to 1896 was postulated recently on the finding of three ancient bone fragments, believed to be analogous to *N. forsteri* bone structures, in a cave located adjacent the Brisbane River. These bone samples were carbon dated at 3,500 years before present (Kemp and Huynen, 2014), but were unable to be confirmed with DNA as being *N. forsteri.*

Early efforts to use genetic methods to answer this question of endemic vs translocated populations often yielded ambiguous results (Frentiu et al., 2001). Despite improvements in molecular markers, genetic data have provided variable levels of confidence to assess the status, endemic or translocated, of the Brisbane River and North Pine River populations. It is important to briefly summarise the results and the conclusions of these studies to highlight that, in this specific case, but also in general for genetic studies more broadly, different translocation hypotheses are challenging to distinguish using descriptive methods alone, even when multiple genetic markers are available.

Frentiu et al. (2001) supported the translocation hypothesis based on one individual from the Brisbane River sharing a rare mtDNA haplotype, not found in the Burnett River, with two individuals sampled in the Mary River. But the most common of the eight haplotypes found in this study was shared among the three rivers. In addition, rare haplotypes are often missed in small size samples, and only 13 or 14 individuals from each river were analysed by Frentiu et al. (2001). Lissone (2003) and Lissone et al. (2004) later found that one polymorphic fingerprinting band was private in the Brisbane River, and even if the other eight bands were shared with other rivers, concluded that translocation could be excluded.

Most recently, genetic variation was analysed with modern markers, allowing a finer comparison among genetic pools across the *N. forsteri* range. Eleven microsatellites identified four major genetic pools corresponding to the Mary River and Brisbane River, Burnett River, North Pine River and Tinana Creek, the latter being a tributary of the Mary River (Hughes et al., 2015). The similarity between Mary River and Brisbane River populations was considered a signature supporting the translocation origin of the Brisbane River population, and the distinctiveness of the North Pine River population was attributed to a founder effect resulting from the small number of individuals reported to have been translocated into this system. However, genetic bottleneck signals expected from a recent origin of these two populations were not found, and very similar levels of diversity in terms of heterozygosity and allelic richness were found across all sites. Complete mitochondrial genomes from 71 lungfish (Bishop et al., 2018) found 173 segregating sites in 16,573 base pairs, and 38 out of 41 haplotypes were private in each of the five rivers analysed in the microsatellite study. The three shared haplotypes, however, were shared among the Mary River and North Pine River (two haplotypes) and the Mary River and Brisbane River (one haplotype). These results were interpreted as evidence of the translocation origin of the North Pine River and Brisbane River populations, with the many private haplotypes resulting from new mutations after the translocation and/or the small sample size, but the possibility of an endemic origin was not excluded (Bishop et al., 2018). Finally, a panel of 15,201 SNPs supported a clear genetic divergence between Mary River, Burnett River, and Brisbane River populations, with the latter showing slightly higher affinity with the Mary River compared to the Burnett River and a slightly lower genetic variation compared to both (Schmidt et al., 2018). A possible impact of the O’Connor Mary-to-Brisbane translocations in explaining these findings was speculated.

Here we used three partially published data sets from mitochondrial genomes (Bishop et al., 2018), 11 microsatellites (Hughes et al., 2015), and 5,196 SNPs (Schmidt et al., 2018; unpublished data) to model the possible origin scenarios for the current Brisbane River and North Pine River populations. We applied a model-based approach of Random Forest Approximate Bayesian Computation, both on single marker types and in a combined marker dataset, and we also analysed the different power of these markers to reconstruct simulated translocation histories. Our results provide evidence that the Brisbane River and North Pine River lungfish populations have a genomic signature that indicates each of these rivers contains an endemic component, which should be considered in future conservation management. We also provide information on the power of the ABC approach and of different markers to address questions on past translocation events.

## MATERIAL AND METHODS

### The genetic data set

The genetic data we analysed refer to five lungfish populations located in Queensland (Australia) (**Figure 1a**: Burnett River (BU), Tinana Creek (TI), Mary River (MA), Brisbane River (BR) and North Pine River (NP)). We analysed three different genetic markers, namely the mitochondrial genome (mtDNA; sample sizes: 71; Bishop et al., 2018), 11 microsatellites loci (STRs; sample sizes: 135; Hughes et al., 2015), and 5,196 SNPs obtained from RAD sequencing (sample sizes: 100; Schmidt et al., 2018; unpublished data). Considering that the SNPs data for the Tinana Creek and North Pine River populations are unpublished, we initially performed a Discriminant Analysis of Principal Components (DAPC; Jombart et al., 2010), using the *adegenet* package (Jombart, 2008) in R, showing graphically the relative genetic distance among individuals (**Figure 2)**. The new data have been generated following the same procedure reported in Schmidt et al. (2018).

**Figure 2.**
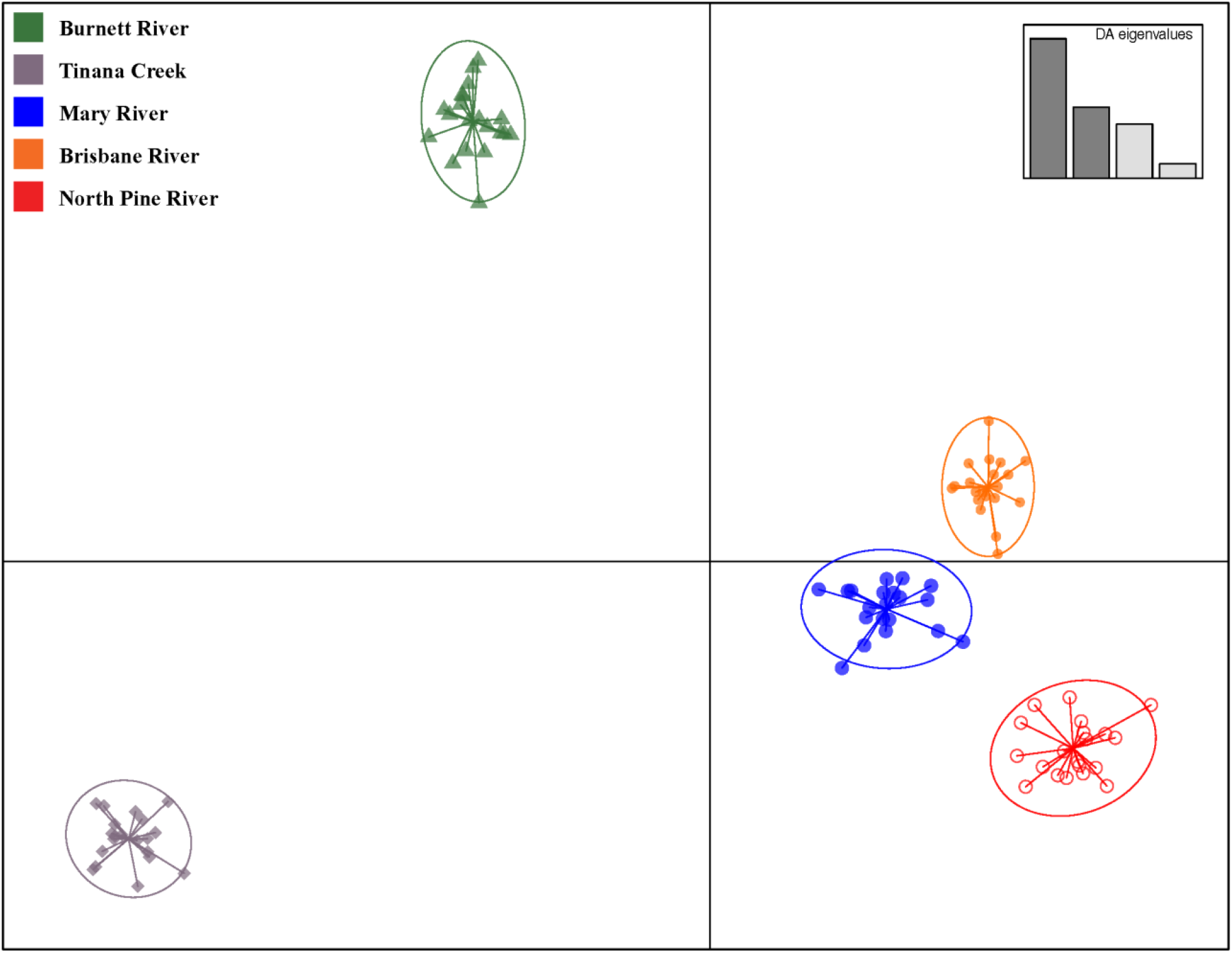
Discriminant analysis of principal components (DAPC) plot showing the relationship of 100 individual lungfish using 5,196 SNPs. Colour coded by sample site.

### The models

Different hypotheses regarding the translocation histories of lungfish populations were tested using a model-based approach. Datasets of variation of the same size of those available were generated by simulation under different evolutionary and translocation scenarios, and the models were scored based on the resemblance between simulated and real data (see below for details). Different models assumed different topologies representing the relationships among populations and different plausible translocation histories. We analysed seven population topologies (labelled Model 1 to 7), each of them assuming five different translocation histories (labelled from -*a* to -*e*) (**Figure S1**).

### Population topologies

Simulations require the definition of the history of the sampled populations. Even if the identification of the best model to describe the relationships between the populations is not of interest in this study, the results on the translocation history would be biased by forcing the populations into a fixed and possibly wrong topological model. We therefore hypothesised seven different relationships (**Figure S1**) based on different logical criteria and let the data identify the best topology that can be used to study the translocation scenarios.

The topology in Models 1- should be considered as a sort of topological null hypothesis, with similar divergence between all pairs of endemic populations. Models 2- assumes a tighter relationship between Tinana Creek and Mary River since they belong to the same catchment. Models 3- add to Models 2- a tighter relationship among rivers having closer estuaries, considering the possibility of gene flow within Northern and Southern catchments during glaciations. Models 4- and 5- are similar to Models 3-, with a longer (or much longer) isolation of Tinana Creek population, which was shown to be highly differentiated at the genetic level (Hughes et al., 2015). Models 6- are designed as Models 4-, but Mary River population is more closely related to the Southern group where Mary River individuals were introduced. Finally, models 7- are similar to Models 6-, but with longer isolation of North Pine River population.

### Translocation scenarios

Models -*a* refer to the null hypothesis for the translocations, i.e., these models exclude recent translocation events and therefore assume that both Brisbane River and North Pine River populations have an ancient origin (as **in Figure 1b**, left panel; see also **Figure S1**). Models *-b* and *-c* assume that Brisbane River and North Pine River populations originated only in recent times, from a translocation event from Mary River (models -*b*; see **Figure 1b**, central panel, and **Figure S1**) or from both Mary and Burnett rivers (models -*c*; see **Figure 1b**, right panel, and **Figure S1**). Under models -*d* and -*e* the Brisbane River and North Pine River populations are both the result of an admixture event between an endemic component and a more recent introgression due to translocations from Mary River (model -*d*; **Figure 1b**, central panel, and **Figure S1**) or from Mary River and Burnett River (models -*e*; **Figure 1b**, right panel, and **Figure S1**).

### Simulation details and summary statistics

All the parameters of the models are listed in the **Table S1** together with their associated prior distributions. Historical evidence was used to define the prior distributions of the parameters related to the translocation events. For the parameters related to topologies (divergence times and population sizes), large uniform (or loguniform) distributions were used. We assumed that endemic populations have constant population size to avoid over-parametrization of the models. For each model we drew 50,000 combinations of demographic parameters from prior distributions and generated a simulated dataset using the *fastsimcoal2* backward (coalescent) simulator (Excoffier et al., 2013). We separately generated mitochondrial, STR and SNP data, considering the different effective population sizes at nuclear and mitochondrial markers. We used a log uniform prior for the mitochondrial mutation rate, between 0.011 and 0.125 mutations per nucleotide per million years, as in Burridge et al. (2008). When simulating STR loci we considered a generalised stepwise mutation model (GSM; Estoup et al., 2002), with average mutation rate across loci having a normal prior distribution with mean equal to 0.0005 and variance equal to 0.0002. At each STR locus, the mutation rate had a gamma distribution with shape parameter equal to 2, as in Marino et al. (2013) (**Table S1**). The simulated data have been summarised by different summary statistics. For the mtDNA we calculated the number of haplotypes, the haplotype diversity, the total and private number of segregating sites, the average number of pairwise differences for each population and Tajima’s D, the global and pairwise F_ST_ and the mean number of differences between pairs of populations. For the STRs, we calculated the mean and standard deviation over loci of the number of alleles, of the heterozygosity, of the modified Garza-Williamson index for each population, and the global and pairwise F_ST_. For the SNPs we calculated the heterozygosity, the FST, and three categories of segregating sites: per population, private, and fixed for opposite sites. We calculated the summary statistics using *arlsumstat* (Excoffier and Lischer 2010) and the three categories of segregating sites through a home-made R script.

### Strategy of model comparison through ABC-RF

The model comparison was performed through an ABC Random Forest (ABC-RF) approach (Pudlo et al., 2015). The ABC-RF, compared to the classical ABC algorithm (Beaumont et al., 2002) works well also with a reduced number of simulations per model and an increased number of summary statistics (Blum & Francois, 2010), thus allowing the study of several complex models. With ABC-RF, the classifier is constructed from simulations from the prior distribution for all the models that must be compared (called reference table) via a machine learning RF algorithm. Once the classifier is constructed and applied to the observed data, the posterior probability of the resulting model can be approximated through another RF that regresses the selection error over the statistics used to summarize the data. Reliable estimates of the posterior probabilities can be achieved with just a few thousand simulations, and the informative statistics are systematically extracted from the pool used to summarize the data (Pudlo et al., 2015). The ABC-RF model selection estimates have been obtained using the function *abcrf* from the R package *abcrf* and employing a forest of 500 classification trees, a number suggested to provide the best trade-off between computational efficiency and statistical precision (Pudlo et al., 2015).

The demographic models detailed in the previous paragraph were compared exploiting a hierarchical approach. We initially identified, within each of the seven topologies tested, the translocation dynamic showing the better fit with the data, thus performing seven independent model selection experiments (e.g., comparing *1a* to 1*e*, then comparing *2a* to 2*e*, etc.). For models 1 and 2 we performed a comparison among 5 models, while the specificity of the other models required a comparison among 8 (models 5) or 11 (models 3, 4, 6, 7) models. This difference is due to the different order of the branching events that requires specific tests. Models were compared four times: separately considering the three types of genetic markers (mitochondrial genomes, STRs and SNPs), and combining them in the same reference table. After this first run of comparisons, we explicitly compared the best scenarios emerging from each topology for each genetic marker considered, and for the whole dataset comprising all the genetic markers together, to identify the model best representing the observed variation, both for the topology and the translocation dynamic.

The confusion matrices and the out-of-bag classification error (CE) were computed in each comparison. The CE measures the proportion of simulated data where the true demographic history is not correctly identified by the classification algorithm. CE is also informative about the respective inferential power of the different genetic markers used. To identify which marker or combination of markers may lead to the best classification algorithm, we also evaluated the Prior Error Rate, which is the average value of the misclassification errors (CE) computed for each scenario (thus assuming an identical prior probability for each model).

The variation generated by the different demographic models, and the position of the observed data, were visualised through a Linear Discriminant Analysis (LDA).

### Parameter Estimation

The parameter values of the most supported model were estimated after increasing the number of simulations to 500,000 and by maximizing the fit between observed and simulated data. One percent of the simulations closest to the observed dataset was retained for the estimation, after a *logtan* transformation of the parameters (Hamilton et al., 2005) and using the local linear regression approach (Beaumont et al., 2002). We estimated parameters considering all the genetic markers together in the same analysis. As suggested by Wegmann & Excoffier (2009), we linear-transformed the vector of summary statistics into partial least square (PLS) components through the *findPLS.r* script within the ABCtoolbox package (Wegmann et al., 2010).

### Additional Simulations to Test for Robustness of Parameter Estimates

We performed an additional set of simulations to analyse in more detail the accuracy of our procedure in estimating low levels of translocation from different sources. To do this we generated 100 pseudo-observed datasets (pods) under known conditions: a fixed topology (Model-*1e*), the effective population sizes and the divergence time fixed to the mode values previously estimated for the selected model, and all the four translocation parameters fixed to a proportion of 0.01. We performed 100 estimations of the proportions of translocated individuals through the same procedure used to analyse real data and assessed the quality of the estimates by computing four indices: the bias, the root mean square error, the factor 2 and the 95% coverage. The bias was calculated as 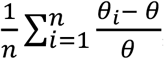, where *θ_i_* is the estimator of the parameter *θ* (true value), and n is the number of pods used (100 in our case). Because bias is relative, a value of 1 corresponds to a bias equal to 100% of the true value.

The root mean square error (RMSE), is calculated as 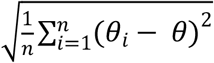. The factor 2 corresponds to the proportion of the estimated median values within the interval defined by the 50% and the 200% of the true parameter value. Finally, the 95% coverage is the proportion of times that the known value lies within the 95% credible interval of the estimates.

## RESULTS

### Comparison of models

**Tables S2** to **S8** report the detailed results for the first hierarchical step, i.e., the comparisons between models that assume the same topology. For example, in **Table S2**, we compared all models with the same “null” topology (equal differences between all pairs of endemic populations, models 1-), but with different translocation scenarios (model 1a to 1e). The comparison within the same topological hypothesis requires more than 5 alternatives (-*a* to -*e*) in some cases, since different branching orders require independent treatment in the simulations. For example, for models 3a, the split between Brisbane River and North Pine River can be simulated as the first, the intermediate, or the most recent in the topology (models 3a_1, 3a_2, and 3a_3 in **Table S4**, respectively).

In general, the models showed a good identifiability, with a Classification Error (CE, measuring the fraction of data simulated with a certain model not classified as originating from that model) in many cases below 10% and in some cases lower than 0.5% (**Figure 3**; **Tables S2-S7**; **Figures S2-S3**). The highest CE values are observed in comparisons accounting for similar models (e.g., within Models 4, 6 or 7), whose specific topology definition made it essential to test eleven different models to explore the different demographic histories proposed. Some differences in CE levels also emerged among markers, with mtDNA and STRs showing higher Classification Errors than SNPs (**Figure 3**; **Figure S2-S3**).

**Figure 3.**
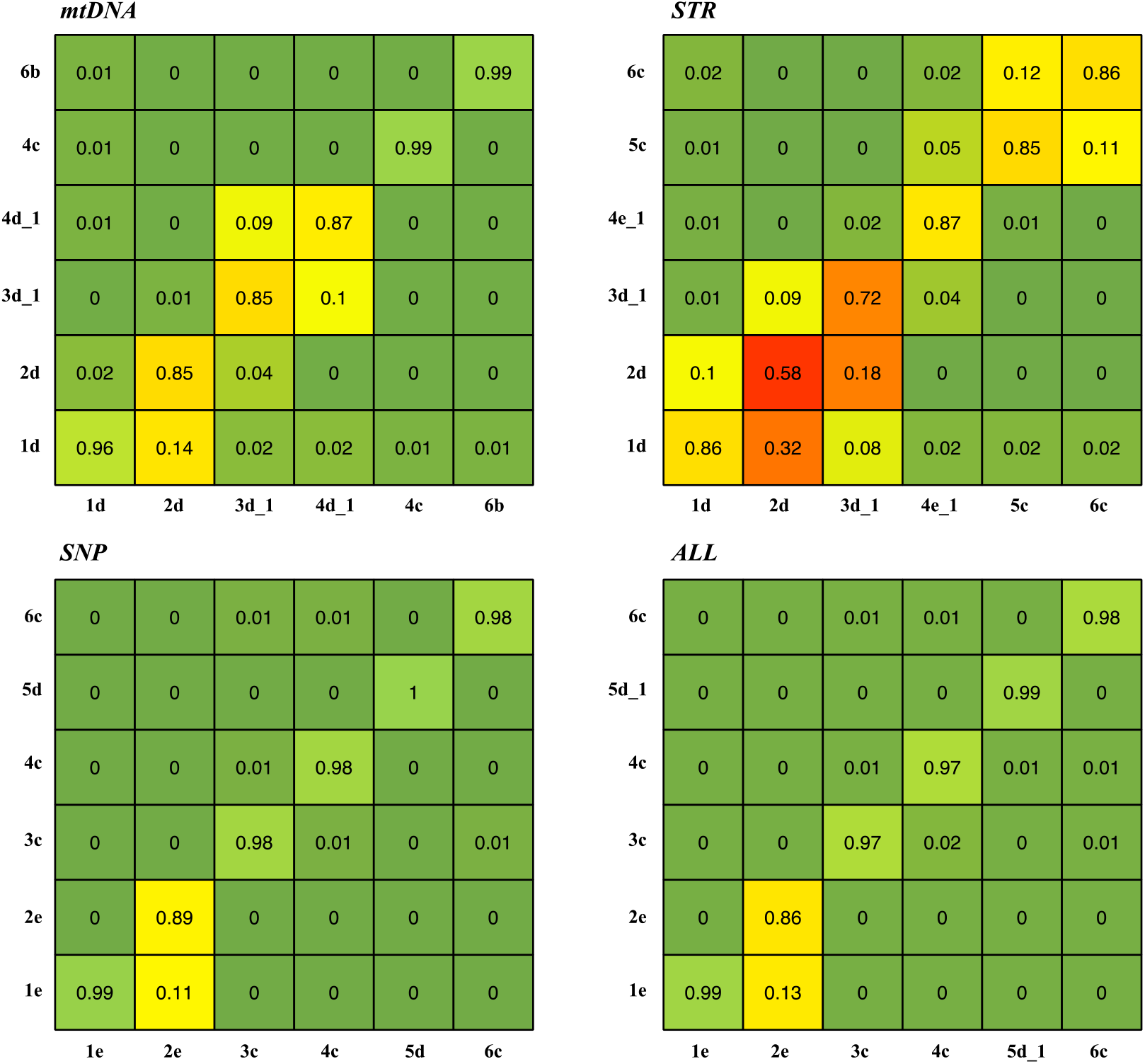
Confusion matrices (for each marker and overall) obtained when the most supported models for each topology are simulated. Lines refer to the simulated model, and columns refer to the fraction of simulations supporting each model. Colour codes are inverted for diagonals (the correct model is recovered) and all the other cells (the wrong model is recovered), to visualize with the same colours problematic inferences.

This is also confirmed by the estimated prior error rates (**Tables S2-S12**) supporting the view that the simulated SNPs dataset is more informative. STR statistics always produce the highest values of prior error rate, about 10 or 20 times higher than those resulting from SNP data in the same model comparison (**Figure S3**).

When applied to real data, the models without translocations (models -*a*) were never selected in the single marker data sets, the global data set, or in the comparisons within the same topology (**Figure 4**; **Tables S2-S12**). Model -*b* (translocated origin of Brisbane and North Pine River populations from Mary River individuals) receives the support only from the mtDNA data set and only under topologies 6 and 7. The most informative dataset (all markers jointly analysed) supports the translocation scenarios -*c*, -*d*, or -*e* when different topologies are considered.

**Figure 4.**
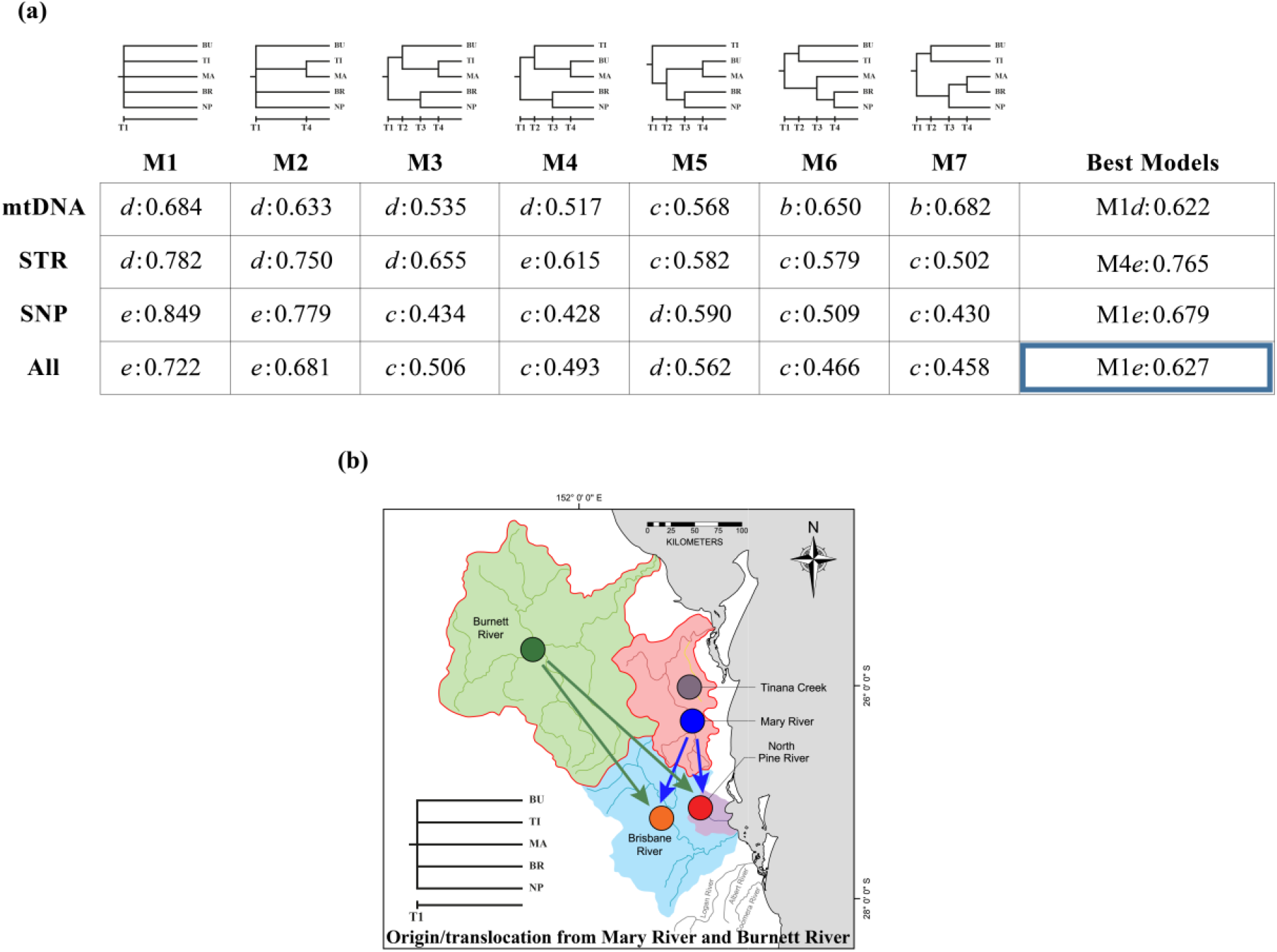
a) Results of the hierarchical model selection procedure. Each column (from M1 to M7) shows the model best accounting for the observed variation within each of the seven topologies represented above the Table. Within each cell is reported the most supported translocation scenario for that topology, with its posterior probability (-*a*: translocation origin is excluded; -*b* and -*c*: BR and NP populations have a translocation origin; -*d* and -*e*: mixed model; see **Figure S1** for details). The last column shows the results of the comparison among the best models resulting from each topology for each genetic marker considered. The highlighted cell represents the model used to estimate parameters. b) Schematic representation of the topology and demographic dynamics of the most supported model considering all the genetic markers together. BU = Burnett River; TI = Tinana Creek; MA = Mary River; BR = Brisbane River; NP = North Pine River.

Once the model best accounting for the observed variation within each topology was identified, we performed a second step of selection, comparing the models supported in the previous step by the whole data set (all markers). Models 6 and 7, in their scenario c which produced the best fit in the previous step (the scenario with Brisbane River and North Pine River that entirely descend from translocated individuals) are identical and six (instead of seven) models were therefore compared (see **Figure S1**). In general, the ability of the classifier to detect the correct model in this comparison was large, with CE between 0.5% and 13% for the SNPs or the complete data sets (**Table S9**). When relying on STRs only, the CE increases considerably, reaching values close to 40%.

The complete data-set favours model 1e, i.e., the simplest topology with all populations equally differentiated, and a mixed origin of Brisbane River and North Pine River from endemic ancestors and individuals introduced from both Mary and Burnett rivers (**Figure 4**). The same topology was also supported by the mtDNA and SNP dataset when separately analysed, and the same translocation history was supported by the STR and SNP dataset, when separately analysed. The posterior probabilities associated to these models range between 62% and 76% depending on the genetic marker considered, four times higher than that expected a priori. We are therefore confident that model 1e is the best model explaining the genetic variation of the lungfish populations we analysed.

We performed an LDA on simulations from the best models (identified for each topology) considering a set of statistics from all the genetic markers (**Figure S4-S5**). The variation generated by these models overlaps eventually along the first three LDA axes, but the projection of the simulations generated under the model selected shows that the observed data are well captured by Model 1e (**Figure S4-S5)**.

### Parameter Estimation

The parameters of the model 1e were estimated using the complete data set and the 30 first PLS components to summarize the data (**Table 1**; **Figure S6**). All but one of the R^2^ values (representing the fraction of the total variance explained by the PLS components used) are between 34% and 75%, indicating that parameter estimates can be considered reliable (Neuenschwander et al., 2008). As expected, the translocation age, an almost fixed parameter with a priori distribution ranging from 1 to 5 generations, cannot be estimated.

**Table 1.** Point estimates (Median and Mode values), the 95% high posterior density (HPD-Lower and Upper bound), and the coefficient of determination (R^2^) obtained for each parameter. Px-y represents the proportion of translocated individuals from x to y. Ttr is the translocation time, Nx the effective population size of the population x, T1 the simultaneous divergence time of the five populations as modeled in the selected topology.

| | Median | Mode | 95% HPD-LowB | 95% HPD-UppB | $R^2$ |
| --- | --- | --- | --- | --- | --- |
| **$P_{BU-NP}$** | 0.0856 | 0.0612 | 0.0000 | 0.2498 | 0.67 |
| **$P_{MA-NP}$** | 0.3048 | 0.1998 | 0.0000 | 0.8121 | 0.40 |
| **$P_{BU-BR}$** | 0.3273 | 0.2664 | 0.0193 | 0.7394 | 0.64 |
| **$P_{MA-BR}$** | 0.7949 | 0.8972 | 0.2989 | 1.0000 | 0.36 |
| **$T_{tr}$** | 4.1845 | 4.5329 | 1.0000 | 1.6877 | 0.03 |
| **$N_{BU}$** | 4,076 | 2,760 | 549 | 11,806 | 0.76 |
| **$N_{MA}$** | 4,743 | 3,368 | 824 | 11,998 | 0.76 |
| **$N_{NP}$** | 278 | 208 | 200 | 727 | 0.38 |
| **$N_{BR}$** | 356 | 200 | 200 | 2,640 | 0.34 |
| **$N_{TI}$** | 2,203 | 1,483 | 210 | 8,051 | 0.75 |
| **$T1$** | 820 | 630 | 300 | 1,949 | 0.54 |

We estimated three sets of parameters: the effective population sizes, the divergence time, and the proportion of translocated individuals from Mary and Burnett Rivers to Brisbane and North Pine Rivers. The five populations show different effective sizes, with higher values for Mary River, Burnett River and Tinana Creek (some thousand of individuals) and lower values for North Pine and Brisbane Rivers (some hundreds of individuals). The simultaneous divergence between all five populations (assumed by the most supported topology) was estimated at 600-800 generations ago, i.e., between 15,000 and 26,400 years ago assuming three to four lungfish generations in 100 years (Schmidt et al., 2018). The estimated fraction of Brisbane individuals that are estimated to have had a Mary River or Burnett River origin at the translocation event is high (**Table 1**; **Figure S6**). The values with the highest probability for these two source rivers are approximately 90% and 27%, respectively, with median values slightly different (79% and 33%, respectively). Assuming either the values with the highest probability or the median values, the total fraction of endemic components in the Brisbane River is estimated between 7% and 14%. The same fractions estimated for the North Pine River indicates a smaller impact of the translocations (**Table 1**; **Figure S6**), with a fraction of endemic component between 65% and 75%.

### Additional simulations to analyse the accuracy in estimating endemic and introduced ancestry

This set of simulations was motivated by the large fractions of introduced ancestry estimated especially in the Brisbane River and considering the fact that historical records indicate that the number of introduced individuals was low. The rationale of the question addressed is to identify intrinsic factors of the methods we implemented that may have produced an over-estimation of the introduced genomic components.

Quality indices were calculated to detect potential low proportions of translocation if present in our observed dataset (**Table S13**). The 95% Coverage values indicated that, for the four parameters analysed, the true value always falls within the 95% confidence interval of the estimates. However, the other indices showed a worse situation, in particular the bias, always showing an overestimation (of about ten times in case of the proportion of translocation from Burnett River to Brisbane River), and the Factor 2, that in all the four cases is zero (Pnb and Psb), or almost zero (Pnm and Psm). It is however worth noting that, being the simulated translocation proportion very small (0.01), any estimated proportion larger than 0.02 would not contribute to the Factor 2 statistic.

## DISCUSSION

Our reconstruction of the Australian lungfish translocation history suggests that “One of the most famous things in Australia” (Longman, 1925) was originally distributed in all the main rivers where it is found today. In particular, the current Brisbane River genetic pool appears to be largely descendent from a few individuals collected mainly in the Mary River and released by the bank manager Daniel O’Connor more than a century ago, as a documented translocation event by the Royal Society of Queensland.

The estimated endemic component comprises approximately 10% of the current gene pool, which is exceptionally low if a pre-existing endemic population existed, indicating the population is now dominated by the genes from these translocated individuals. For this situation to have developed, different non mutually exclusive scenarios could be postulated: (1) the translocated lungfish have far outperformed any pre-existing endemic population, (2) additional undocumented translocation events could have taken place in the last century, or (3) the original endemic Brisbane River population prior to the translocations was very small. In a second river, where the presence of a translocated population is under question, the North Pine River, the fraction of the endemic genes is much larger, around 70%. Historical records suggest that the source of the translocated individuals was also from the Mary River with some possible later but undocumented translocations from the Burnett River. The genetic data suggest a larger contribution (approximately three time larger) of the Mary River gene pool compared to the Burnett River, in both the Brisbane and North Pine rivers.

These proportions, despite the large number of markers of different type, have large confidence intervals and should be interpreted cautiously. The main unanswered question we addressed, however, was about the presence or not in the Brisbane River and the North Pine River of an endemic genetic component, and we believe that our analysis supports a conclusion there is. The best fitting model selected by our hierarchical process of model comparison excludes a complete non-endemic (translocation only) ancestry in these rivers, and this result is found not only when all markers are simultaneously analyzed, but also when they are separately considered. Importantly, the simulated datasets suggest that if the real scenario would have excluded an endemic component (models b and c under different population topologies), our approach would be unlikely to have missed it. Similarly, the models without translocations can be confidently excluded.

Other estimated features and parameters of the best model support a similar genetic divergence (before the translocations) of the five different rivers, estimated at 15 to 26 thousand years ago, and two classes of effective population sizes, smaller (some hundreds) in North Pine River and Brisbane River and larger (some thousand) in Mary River, Burnett River, and Tinana Creek. The first topological inference may reflect a similar divergence due to a real split event related to the last glaciations, but more likely is just telling us that these populations diverged for enough time (possibly also with local adaptation) to reduce the power to infer a branching scheme. In our study we considered the population topology as a nuisance parameter, to be accounted for to improve the comparison between models assuming different translocation patterns, but of no real interest. The effective population sizes we estimated for the five populations are reasonable in both absolute and relative terms, and do not point to an immediate risk of extinction. The smaller effective population size estimated for the Brisbane River is an interesting outcome given it contains a relatively large and comparable catchment size (7,015 km^2^) and environment compared to the Mary River (9,595 km^2^). This result can possibly be explained by a largely translocated origin of the Brisbane River population. On the other hand, considering that the North Pine River is considerably smaller than the others at only 348 km^2^, the smaller effective population size we estimate is expected.

Several genetic markers and statistical methods have been used in recent years to identify historical translocation events. Here we show that an additional modelling step can be used to improve these inferences further. In fact, explicit and detailed demographic models, and the estimation of their probability through simulation, outperform descriptive approaches that require interpretation of the patterns and integration of the results produced by different analyses and different markers. As a simple example of this difference, we note here that the descriptive analyses based on the SNP dataset (see DAPC in **Figure 2**), where Brisbane River and North Pine River individuals shows some degree of resemblance with those sampled in Mary River, is visually compatible with a translocation event, but do not exclude models without translocation or without endemic contribution that were frequently suggested for this species in previous studies. The model comparison approach we applied here combines different sources of genomic data and quantifies probabilities and confidence intervals of possible outcomes.

The main conclusion from this work that there is a high probability that Brisbane River and North Pine River contain a mix of endemic and translocated ancestry has been reached using a combined data set, but also limiting the analysis to the single non-recombining marker with less than 200 segregating sites (mtDNA), to independent and classical multiallelic nuclear markers (STRs), or to >5,000 biallelic random nuclear loci (SNPs) (**Figure 4a**). Nevertheless, the study of the simulated data sets clearly shows that the SNP markers are more informative. For example, considering different population topologies, the fraction of datasets simulated under model -*c* (only non-endemic origin of Brisbane River and North Pine River) erroneously assigned to model -*e* (mixed origin) is always smaller than 1% for the SNP panel, but much larger values are found for the other markers (up to 5% and 10% for mtDNA and STR markers, respectively). The larger error rate found in STRs compared to the mitogenomes is rather unexpected, considering that variable sites in the mtDNA are all linked. This could be an effect of the mutational model used for STRs which was not deeply investigated in this system, or to the low genetic variation observed in these markers (Hughes et al., 2015). We did not investigate the effects on the parameter estimation of using separately the three types of markers, but it is reasonable to expect better estimates and smaller support intervals when nuclear SNPs are considered.

### Translocations and conservation

The aim of translocation plans in modern conservation biology is to create or maintain a viable and self-sustainable population or species, when the endemic population is extinct in the wild, or the extinction risk is estimated to be high or predicted to increase (Weeks et al., 2011). When the translocation implies the rescue of an endemic group (also called restocking), the expected genetic benefits are, in the short-term, a fitness increase due to reduced inbreeding depression and decreased frequency of fixed deleterious mutations, and, in the long term, an increase in genetic variation and thus the evolutionary potential. On the other hand, the introduced genetic pool may have negative effects such as outbreeding depression or if the endemic ancestry is reduced significantly up to a point where the rescue becomes a replacement, and the specific traits and genes of the endangered group disappear. Clearly, Daniel O’Connor and the members of the Royal Society of Queensland were only considering the possible benefits of increasing the distribution of this rare and “important” fish, presumably on the assumption that the Australian lungfish were not present at the time in the Brisbane River and the North Pine River. No records of anyone checking this assumption exist.

After more than a century, the genomic data we analyzed suggests that the lungfish were already distributed in the Brisbane River and the North Pine River, and therefore that O’Connor performed a restocking intervention. The introduced animals were very likely genetically divergent from any assumed endemic population, considering the level of genetic divergence found among the Mary River, the Burnett River and the Tinana Creek that have not experienced translocation events that we are aware of. The original endemic gene pool is largely lost in the Brisbane River, but could be argued to have been maintained in the North Pine River. If the endemic populations in these rivers would have survived without the restocking is difficult to guess, but a major outbreeding depression effect can be excluded considering the current population sizes (Hughes et al., 2015).

The implications of these new findings will inform future genetic management considerations for the species. Currently the Australian lungfish is conserved under national and international conservation legislation. The primary Australian legislation, the Environmental Protection and Biodiversity Conservation Act (EPBC Act, 1999), considers all existing populations of a threatened species, irrespective of endemic or translocated origin, as a significant sub-population for the species as a whole. In this case, the mixed origin, endemic and translocated, of the Brisbane River or North Pine River populations is inconsequential from a conservation perspective while the species as a whole is classified as vulnerable to extinction, in that all sub-populations form an important management unit to sustain the species as a whole. However, considering that the local habitats of the five main lungfish populations are quite different, and genetic divergence is high, we expect some level of local adaptation to these different conditions supporting independent management. And if this is particularly true for the rivers where translocation can be excluded, we believe that also the likely admixed populations in the Brisbane and the North Pine River should be followed with great attention analyzing the genetic divergence and the introgression process at different genes. More practically, the value of the Brisbane River and North Pine River lungfish as back up populations to the endemic populations of the Burnett River, Mary River and Tinana Creek, requires careful consideration from a genetic standpoint. These mixed gene pools certainly need to be managed as separate back-up populations and should not to be mixed with the known endemic populations of the Burnett River, Mary River and Tinana Creek, without careful consideration of the genetic implications.

Finally, small remnant populations of the presumed translocated lungfish from 1896 still exist in a number of other river catchments in the Brisbane area, including the Enoggera Creek, Coomera River and Logan River systems. Their genetic analysis could be of great value to better reconstruct the evolutionary and translocation history of this species, and to plan future interventions.

## AUTHOR CONTRIBUTIONS

SG, JMH and GB designed the study. DJS produced the SNPs data. RB and SG analysed the data. RB, SG and GB wrote the manuscript. All the authors discussed and interpreted the results and contributed to the writing.

## ACKNOWLEDGMENTS

GB, SF, and RB are deeply grateful to Jane Hughes and Dan Smith for their hospitality at the Griffith University, Queensland, and for their friendship. Part of this study was funded by the University of Ferrara. RB was supported by MUR grant 201794ZXTL. JMH and DJS were supported by the Griffith University, Queensland.

## DATA AVAILABILITY STATEMENT

Individual genotype data from RADseq are available on Zenodo (10.5281/zenodo.7646503).

