## Supplementary Figures for "Unravelling the mystery of endemic versus translocated populations of the endangered Australian lungfish (*Neoceratodus forsteri*)"

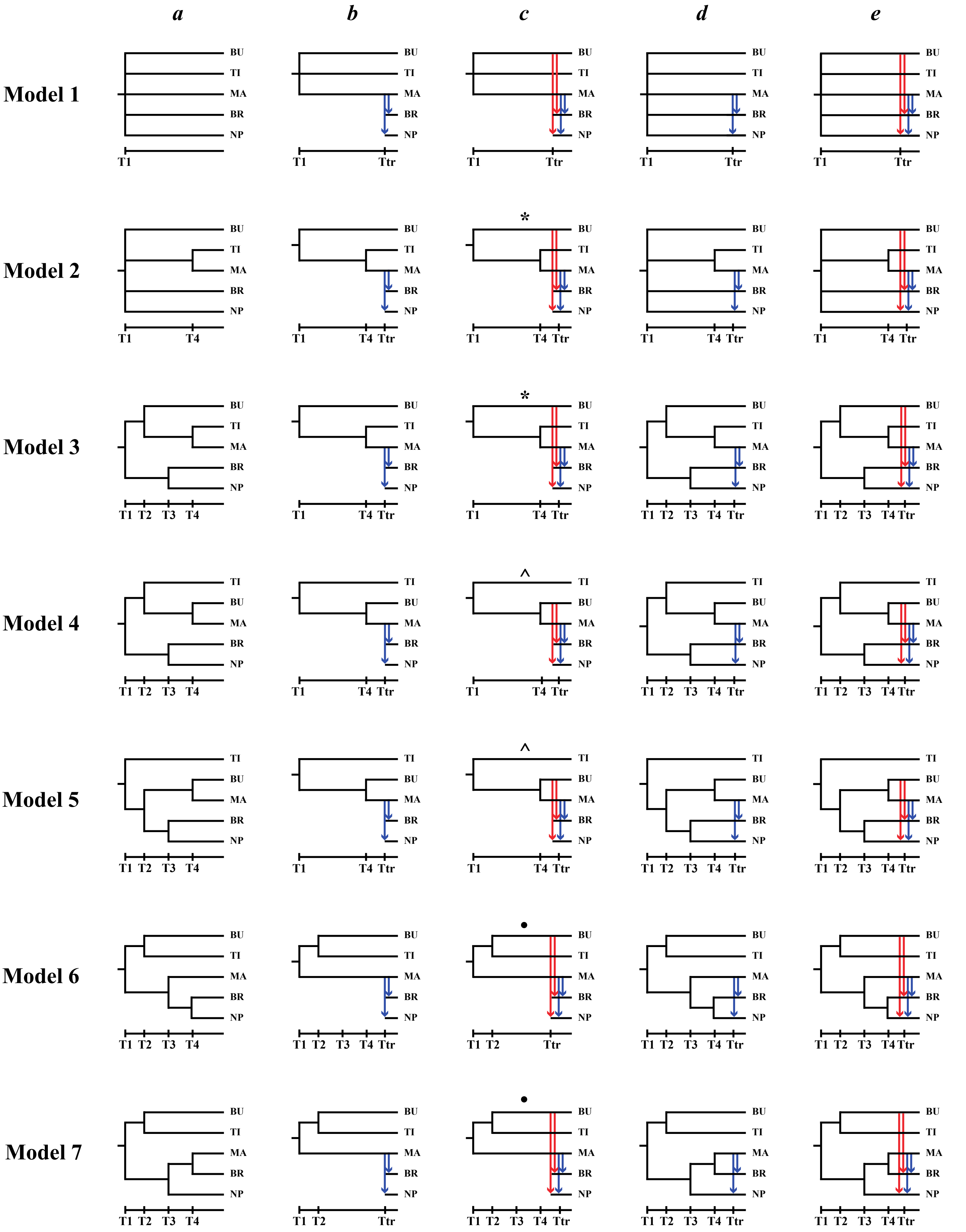


**Figure S1.** Seven population topologies (labelled from Model 1 to Model 7), each of them assuming five different translocation histories (labelled from -*a* to -*e*). Red arrows indicating translocations from Burnett River and blue arrows from Mary River. Pairs of collapsed models, i.e., pairs of models virtually identical, are indicated with symbols (*, ^, ). BU = Burnett River; TI = Tinana Creek; MA = Mary River; BR = Brisbane River; NP = North Pine River.

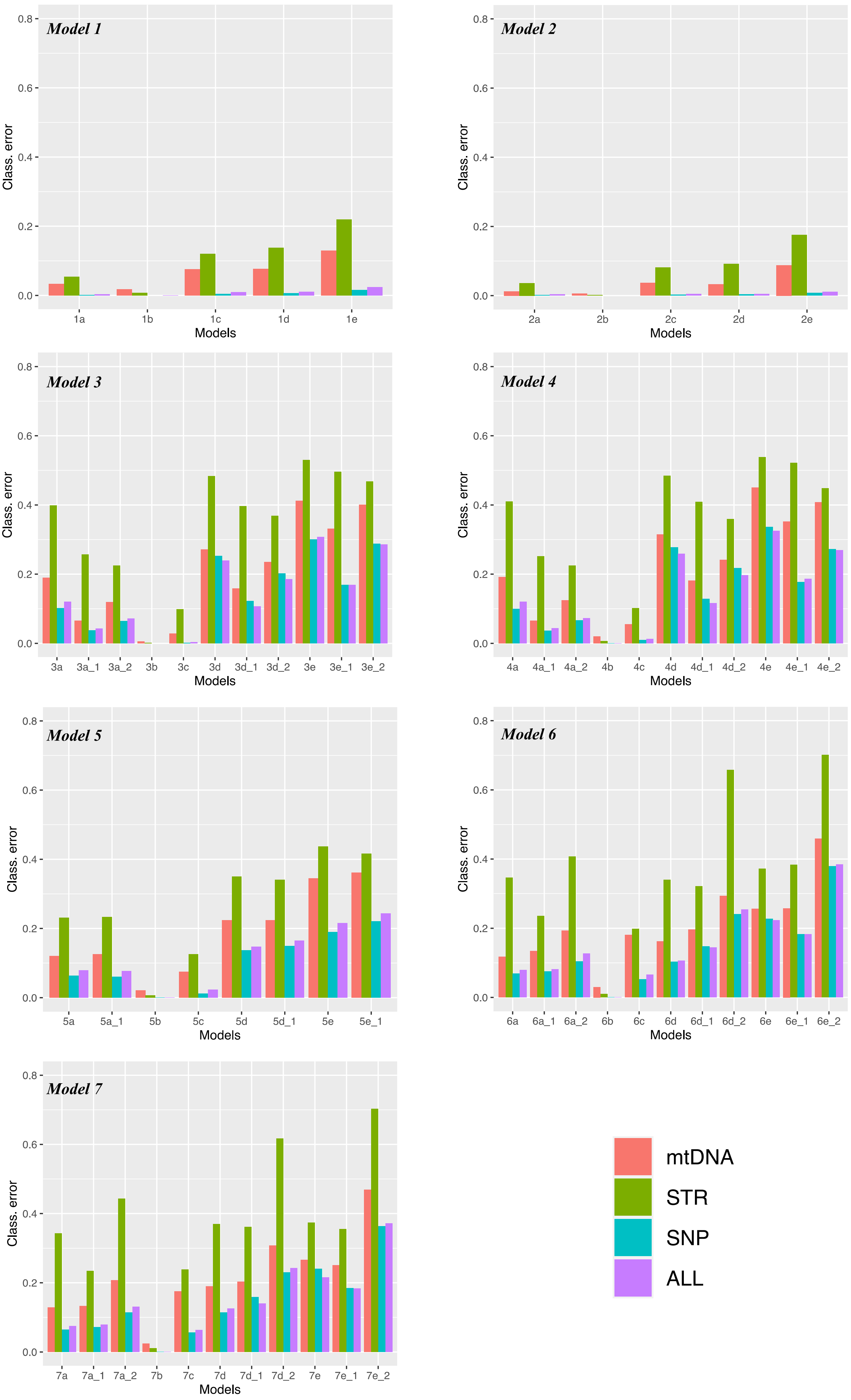


**Figure S2.** Bar plots showing the Classification Errors computed for each comparison within each Model and for each marker.

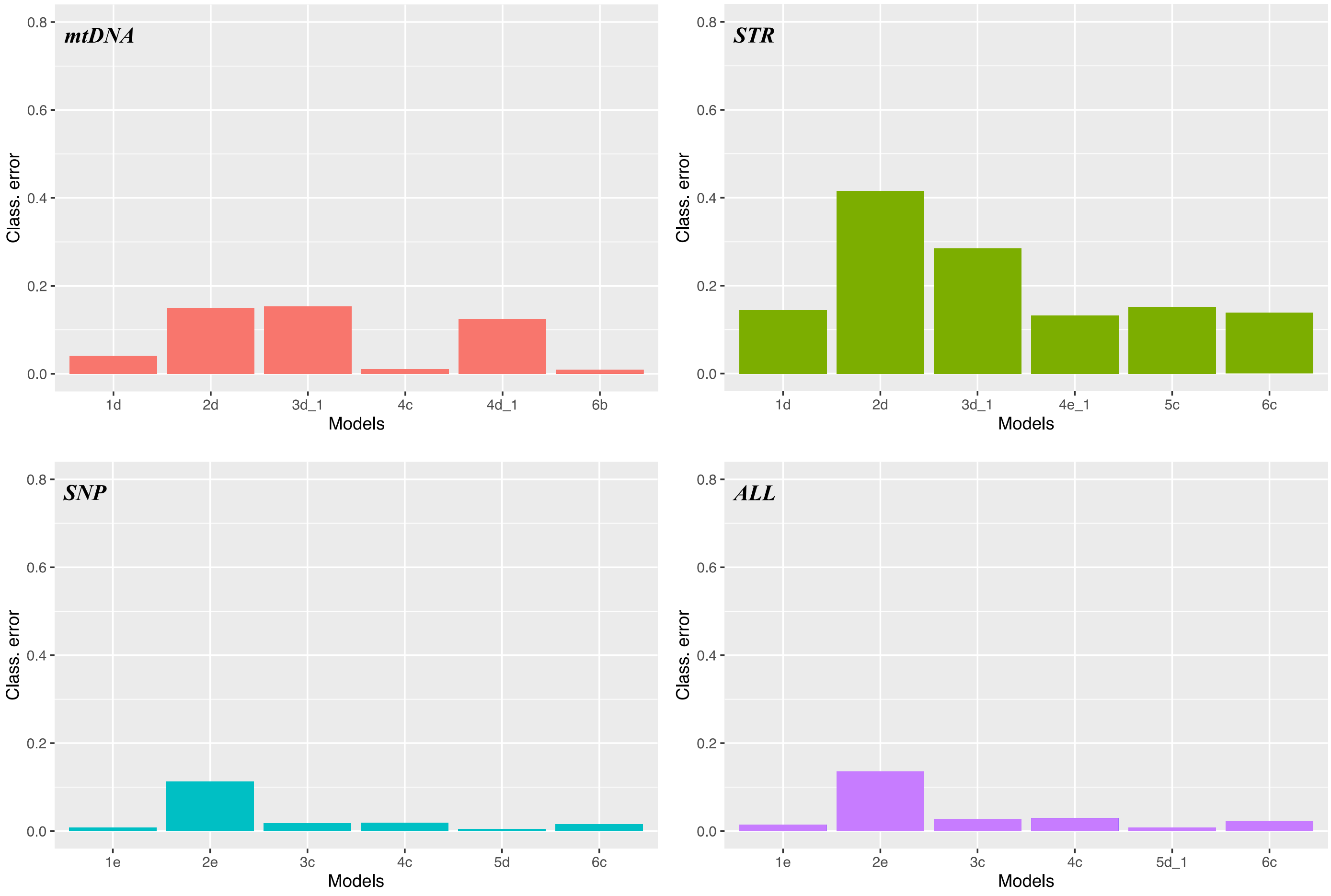


**Figure S3.** Bar plots showing the Classification Errors computed for each comparison among the best models, resulting from each topology for each genetic marker considered.

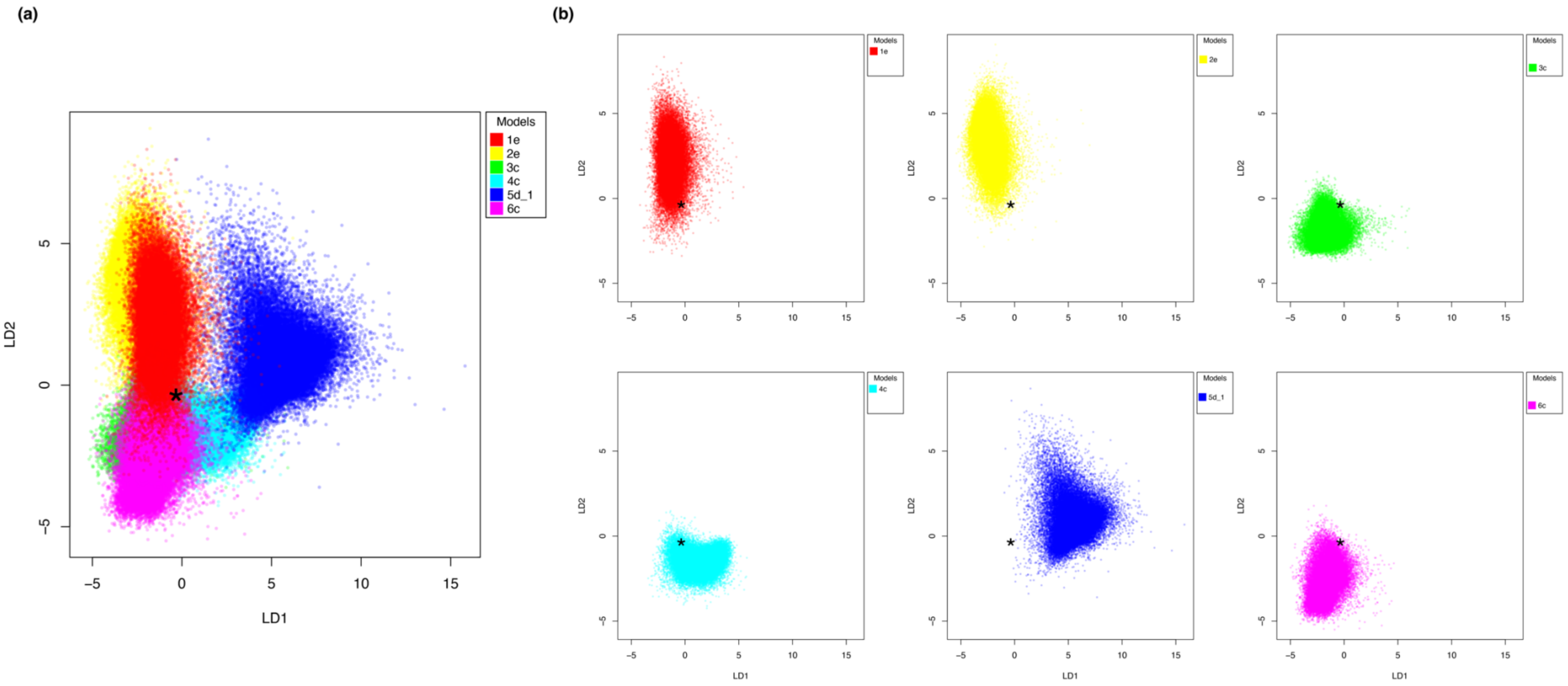


**Figure S4.** a) Linear Discriminant Analysis (LDA) plot of the first and second discriminant functions generated from simulations from the best models (identified for each topology) considering a set of statistics from all the genetic markers. b) Plots showing the distribution of each model separately.

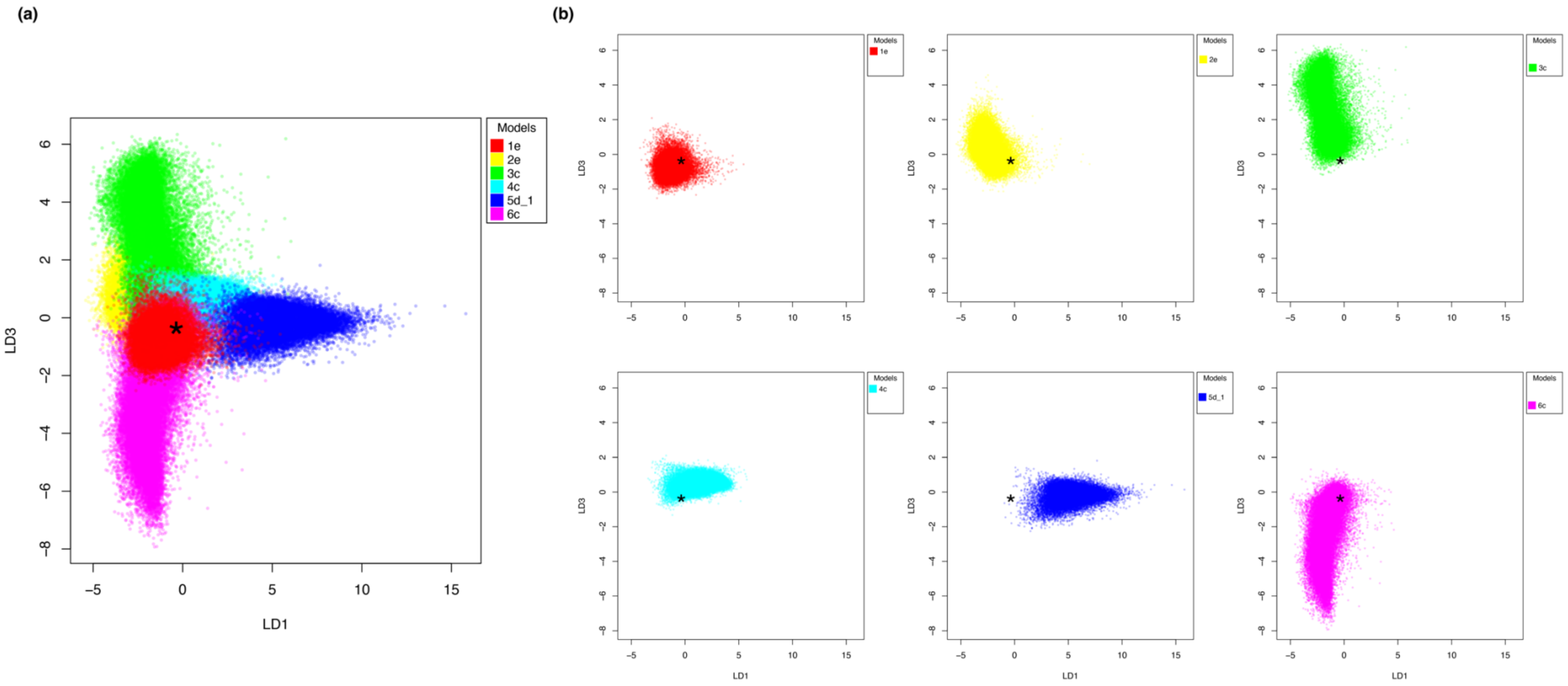


**Figure S5.** a) Linear Discriminant Analysis (LDA) plot of the first and third discriminant functions generated from simulations from the best models (identified for each topology) considering a set of statistics from all the genetic markers. b) Plots showing the distribution of each model separately.

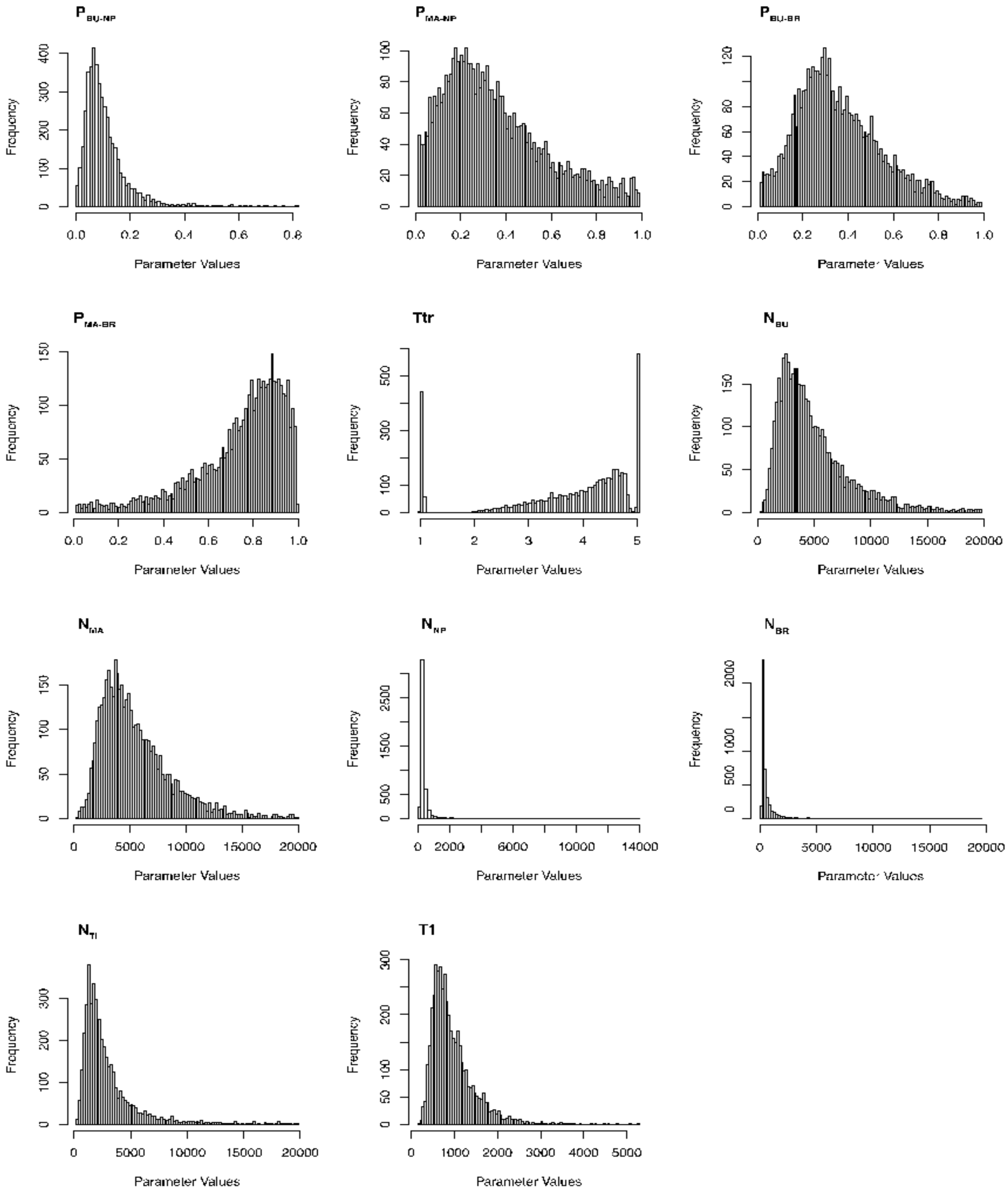


**Figure S6**. Plots of the post distributions of the Models' parameters. $P_{x-y}$ represents the proportion of translocated individuals from x to y. Ttr is the translocation time, $N_x$ the effective population size of the population x, T1 the simultaneous divergence time of the five populations as modelled in the selected topology.
